# An atlas of genetic associations in UK Biobank

**DOI:** 10.1101/176834

**Authors:** Oriol Canela-Xandri, Konrad Rawlik, Albert Tenesa

## Abstract

Genome-wide association studies have revealed many loci contributing to the variation of complex traits, yet the majority of loci that contribute to the heritability of complex traits remain elusive. Large study populations with sufficient statistical power are required to detect the small effect sizes of the yet unidentified genetic variants. However, the analysis of huge cohorts, like UK Biobank, is complicated by incidental structure present when collecting such large cohorts. For instance, UK Biobank comprises 107,162 third degree or closer related participants. Traditionally, GWAS have removed related individuals because they comprised an insignificant proportion of the overall sample size, however, removing related individuals in UK Biobank would entail a substantial loss of power. Furthermore, modelling such structure using linear mixed models is computationally expensive, which requires a computational infrastructure that may not be accessible to all researchers. Here we present an atlas of genetic associations for 118 non-binary and 599 binary traits of 408,455 related and unrelated UK Biobank participants of White-British descent. Results are compiled in a publicly accessible database that allows querying genome-wide association summary results for 623,944 genotyped and HapMap2 imputed SNPs, as well downloading whole GWAS summary statistics for over 30 million imputed SNPs from the Haplotype Reference Consortium panel. Our atlas of associations (GeneATLAS, http://geneatlas.roslin.ed.ac.uk) will help researchers to query UK Biobank results in an easy way without the need to incur in high computational costs.

## INTRODUCTION

Most human traits are complex and influenced by the combined effect of large numbers of small genetic and environmental effects^1^. Genome-wide association studies (GWAS) have identified many genetic variants influencing many complex traits. The largest genetic effects were discovered with modest sample sizes, with researchers subsequently joining efforts to increase the size of the study cohorts, thus allowing them to identify much smaller genetic effects. The UK Biobank^2^, a large prospective epidemiological study comprising approximately 500,000 deeply phenotyped individuals from the United Kingdom, has been genotyped using an array that comprises 847,441 genetic polymorphisms, with a view to identify new genetic variants in a uniformly genotyped and phenotyped cohort of unprecedented size.

The unprecedented size of this cohort has raised a number of analytical challenges. First, storing, managing and analysing the circa 96 million genetic variants for around half a million individuals is, in itself, a substantial endeavour. Second, the collection of samples at this scale has brought up an analytical challenge. As many relatives were unintentionally collected in the cohort, removing them from the analyses as traditionally done in GWAS would entail a substantial loss of statistical power. Third, fitting a Linear Mixed Model (LMM), the standard analytical technique to perform GWAS when the sample contains related individuals, at this scale entails a computational burden which may be beyond the means of many institutions.

The objective of the current study was to perform GWAS for 717 traits in UK Biobank, adjusting for the effect of relatedness to minimise the loss of statistical power whilst avoiding false positives due to family structure, in individuals of White-British descent and to make a searchable atlas of genetic associations in UK Biobank for the benefit of the research community.

## RESULTS

### Data overview

In July 2017, the UK Biobank released genotyped data from circa 490,000 individuals of largely White-British descent genotyped for 805,426 SNPs. We performed GWASs for 599 binary traits and 118 non-binary traits, the latter including continuous traits and traits with multiple ordered categories (Supplementary Table 1). For each of these traits we used a LMM to test 31,415,476 genetic polymorphisms for association. In total, we tested 623,944 genotyped and 30,798,054 imputed genetic polymorphisms imputed using the Haplotype Reference Consortium^3^ as reference panel. All successfully tested polymorphisms are shown in the database and associated downloadable files to allow individual researchers to apply their own quality control thresholds. The summary results presented here are based on quality controlled genotyped polymorphisms (Methods).

The phenotypes selected comprise a mix of baseline measurements (e.g. height), self-reported traits at recruitment (e.g. self-reported depression), and Hospital Episode Statistics (i.e. data collected during hospital admissions) as well as cancer diagnoses from the appropriate UK Cancer Registry. Since UK Biobank is a recently stablished prospective cohort, we allowed for potential differences in statistical power among binary and non-binary traits by splitting the presentation of the data into non-binary and binary traits.

### Heritability Estimates

Heritability estimates inform about the contribution of genetics to the observed phenotypic variation. The heritability of many of the 717 traits analysed here has never been reported, but even if they have been reported it is useful to know how much phenotypic variation is captured by the SNPs in a cohort of the size and interest of UK Biobank. The majority (78.4%) of the traits analyzed had a significant SNP-heritability (P<0.05; Figure 1), with the largest SNP heritability being for ankylosing spondylitis, which was 0.9 on the liability scale. The mean and median heritability among those that were significant were 0.12 and 0.09, respectively. Mean heritabilities were significantly different for binary and non-binary traits (h^2^_Non-binary_=0.16; h^2^_Binary_=0.11; P=1.7x10^-7^). A total of thirty-one traits, all binary, had a heritability estimate close to zero (h^2^_Liability_ < 10^-4^). Only two of those thirty-one traits had no genome-wide significant hits (P<10^-8^) and three of them had more than ten hits. This scenario could arise for monogenic and oligogenic traits for which the model assumptions do not hold or because of false positives. The Manhattan plots for the traits that had the largest numbers of hits are consistent with these hits being false positives and not with the violation of the model assumptions (Supplementary Fig. 1).

**Figure 1.**
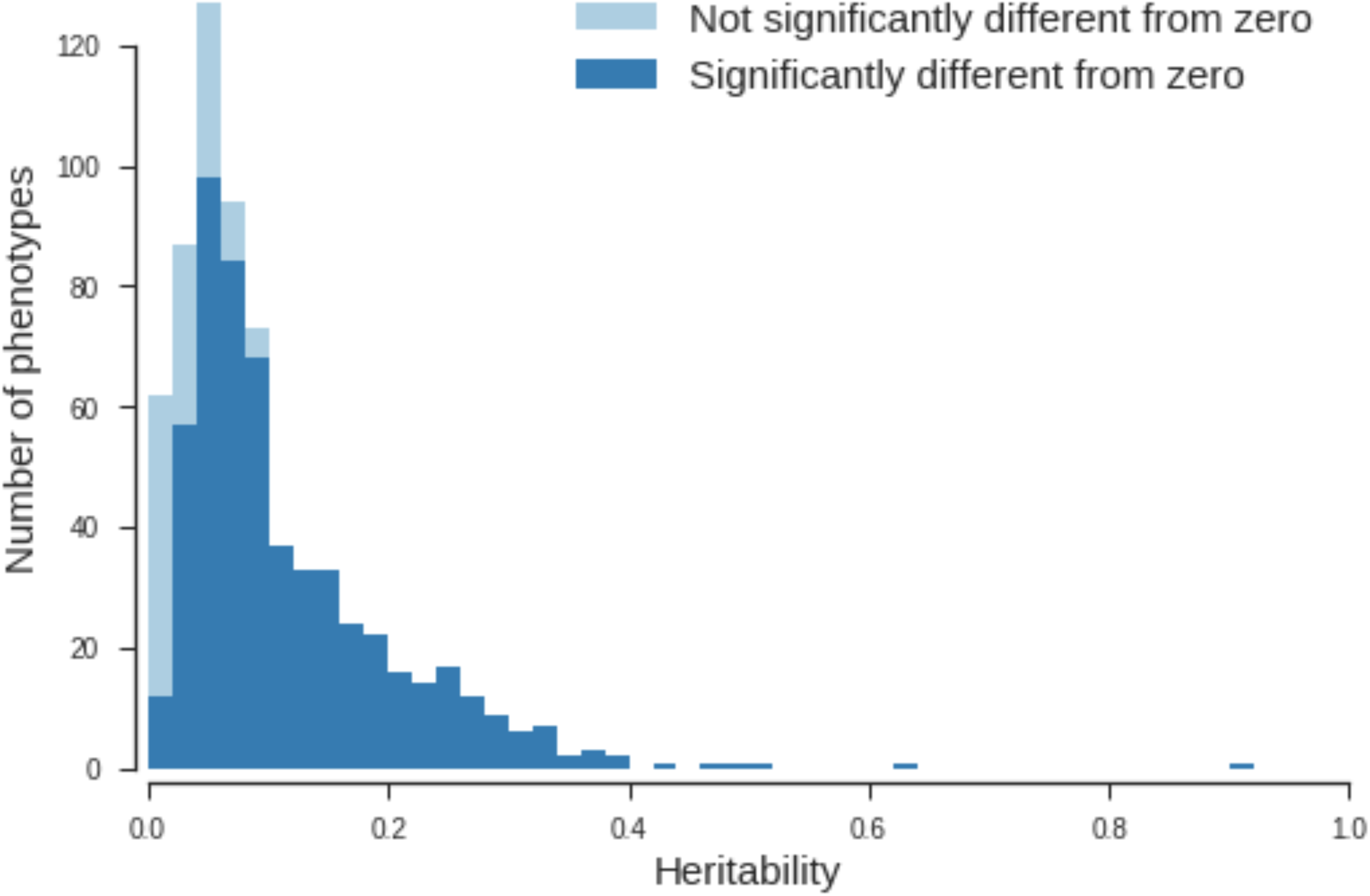
Numbers of phenotypes of different SNP heritability. Colours indicate the fraction of phenotypes with heritability significantly (P < 0.05) different from zero in each bin.

Estimates of genetic and environmental correlations show that for 18% of the pairs of non-binary traits the genetic and environmental correlation changes sign (Supplementary Fig. 2, GeneATLAS web page). Across all non-binary pairs of traits for which the genetic and environmental correlation had the same sign the absolute value of the genetic correlation was smaller in 26% of the cases. Overall, taking into account the size of observed heritabilities, this suggests that the phenotypic covariance of many of these traits is mainly driven by the environment and not genetics (average (cov_g_/cov_e_)–0.23, among traits where cov_g_ and cov_e_ have the same sign).

### Distribution of genotyped GWAS hits among non-binary trait

Just below a quarter of a million of the circa 73 million tests performed across 118 non-binary traits were significant at a conventional genome wide threshold (P<10^-8^) (Supplementary Table 2), and 204,374 were significant after Bonferroni correction (P<0.05/623944*118). The significant associations where distributed across 22,497 leading polymorphisms mapping to 16,148 independent loci (Methods, Figure 2). A large proportion of these associations (37.7%) were within the HLA region (Supplementary Table 2). The fraction of significant polymorphisms was always larger than expected by chance under the assumption that all traits and polymorphisms were independent. For example, even at a threshold of P=10^-2^, we observed a ∼4 fold enrichment of significant associations, consistent with the hypothesis that complex traits are extremely polygenic and determined by many variants of very small effect. In such a case, these effects are so small that we anticipate that it will be practically impossible to detect them at the levels of genome-wide statistical significance currently used as this would require studies of extremely large sample sizes.

**Figure 2.**
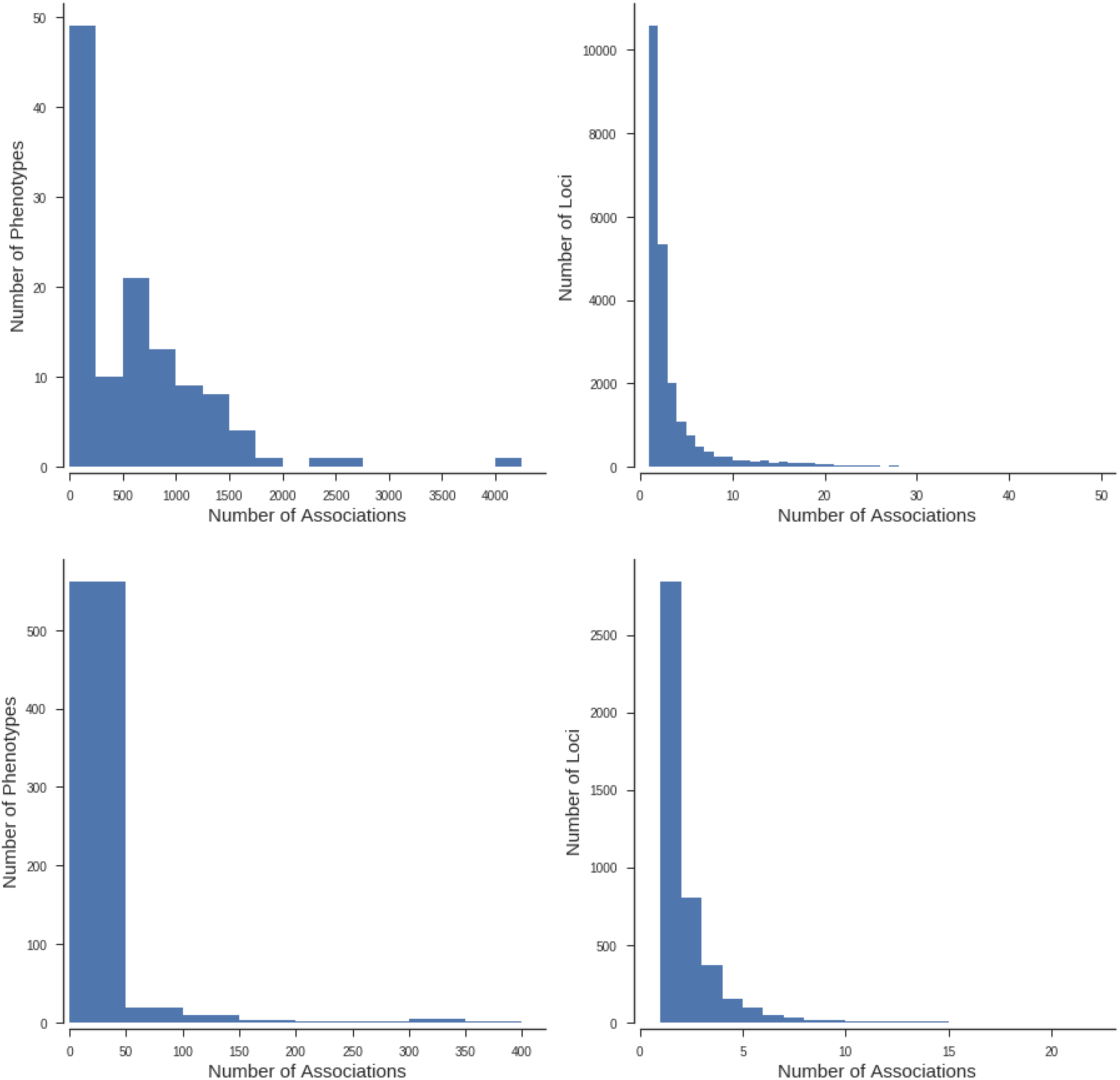
Histograms of numbers of significant associations (P < 10^-8^) for each phenotype (left) and independent locus (right) for non-binary (top) and binary (bottom) phenotypes.

About 6.7% of the tested polymorphisms reached genome-wide significant thresholds (P<10^-8^) for at least one of the 118 tested traits, whilst 78% of the tested polymorphisms were associated with at least one of these 118 traits at a significance level of 10^-2^ (Supplementary Table 3). There were 1405 variants which each were associated with more than 30 of the tested non-binary traits (Figure 3, Supplementary Fig. 3). The intronic variant (rs1421085) within the *FTO* gene had the largest number of genome-wide significant associations, being found to be associated with 57 traits (Supplementary Fig. 3). The variant at the *FTO* locus also had the largest average significance across non-binary traits (P_log-mean_<10^-59^) (Supplementary Fig. 4), which was largely contributed by the associations to anthropometric traits with BMI and Weight measures showing the strongest associations (P<10^-200^). The HLA region contained twenty-nine genetic variants which where significantly (P<10^-8^) associated with 50 or more of the non-binary traits compared to only eleven such variants in the remaining autosomal variants. Four traits ('Standing height', 'Sitting height', 'Platelet count', 'Mean platelet (thrombocyte) volume') had over 6,000 significant associations (P<10^-8^) across 8556 different independent lead genetic variants (Methods). Over three quarters of the non-binary traits had more than 100 genome-wide significant hits distributed in 22,242 different lead genetic variants. Considering the criteria for inclusion of genetic polymorphisms on the genotyping array, the HLA polymorphisms were the most enriched for associations with at least one trait (83% had a P<10^-8^), followed by the Cardiometabolic, ApoE and Autoimmune/Inflammatory criteria, whilst the lowest enrichment was for two low frequency variants categories (“Genome-wide coverage for low frequency variants” and “Rare, possibly disease causing, mutations”). Less than 4 in 100 of these polymorphisms was associated with any non-binary trait (Supplementary Table 4).

**Figure 3.**
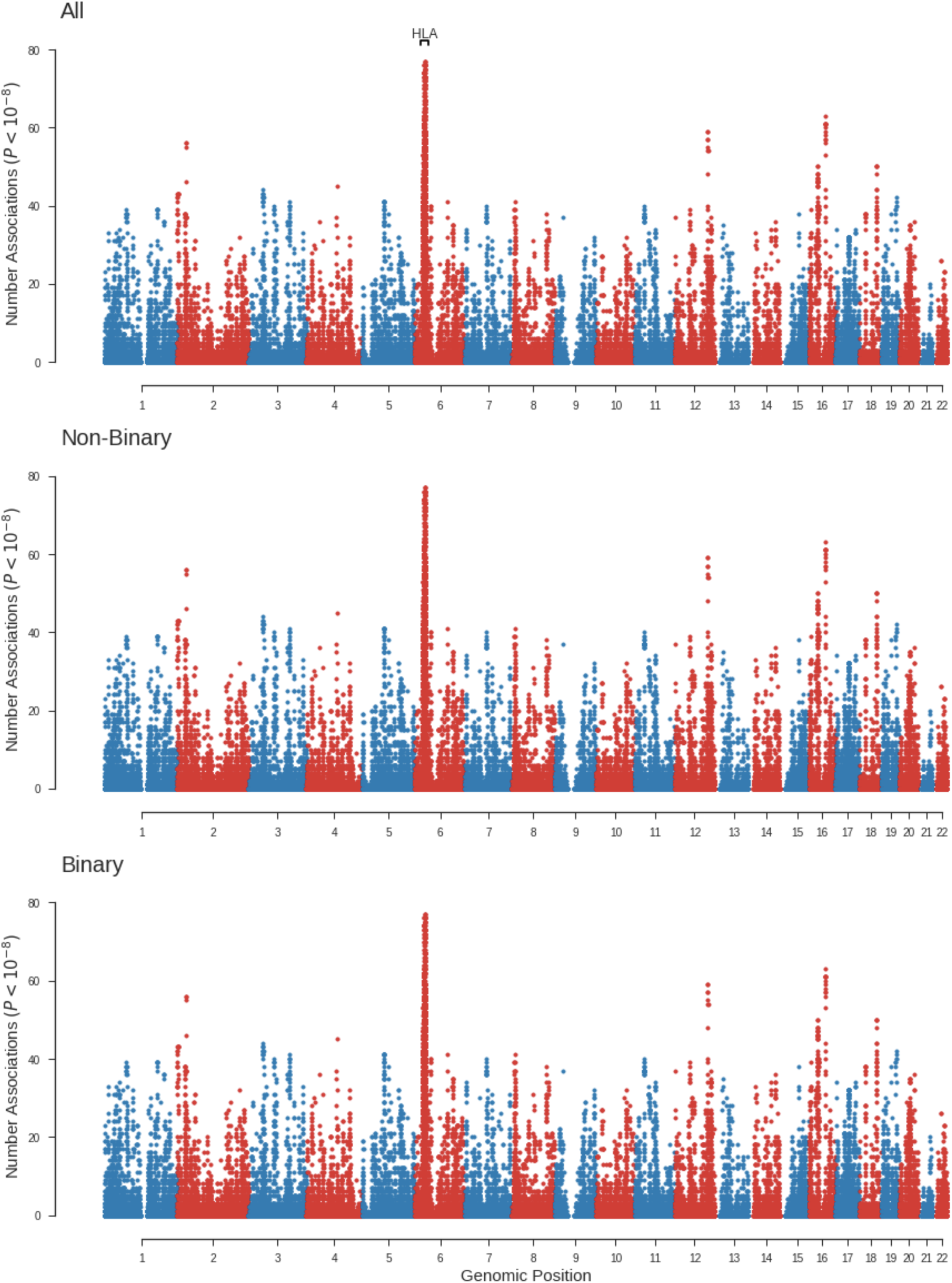
Number of significant associations (P < 10^-8^) at each tested genetic variant for all traits, non-binary and binary phenotypes. The HLA region (±10Mb) is indicated.

We found a significant correlation (r=0.91, P<10^-46^) between the number of hits and the SNP heritability of the traits, suggesting that the number of loci affecting a trait might be proportional to the heritability of the trait (Figure 4, Supplementary Fig. 5). Consistent with this model and variation in the distribution of linkage disequilibrium across the genome, the correlation of the SNP heritability with the number of identified independent loci was similarly high (r=0.89, P<10^-47^). The number of hits (P<10^-8^) per chromosome was highly correlated (r=0.70) with the length of the chromosome covered by the genotyped SNPs (Supplementary Fig. 6, Supplementary Table 5). Although this correlation could arise under a polygenic model where the length of the chromosome is correlated with the number of possible variants affecting the traits, the simplest explanation is that it arises as a consequence of the correlation of chromosomal length and number of variants per chromosome. Comparing the fit of two nested models to explain the number of hits per chromosome as a function of number of genetic variants in the array and length of the chromosome or just the number of genetic variants was consistent with the later explanation (Methods).

**Figure 4.**
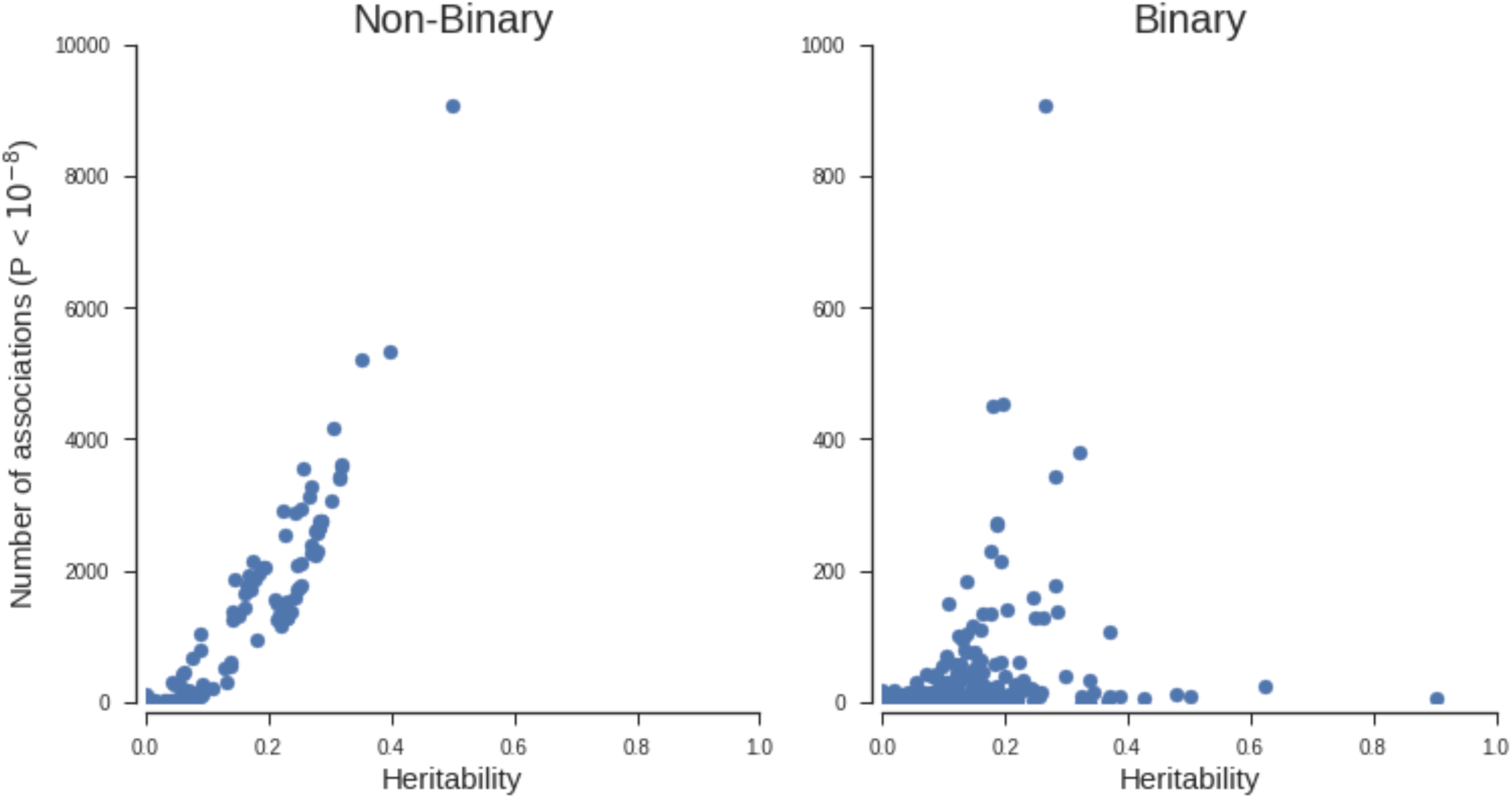
Relationship between estimated SNP heritability and numbers of genome wide significant associations (P < 10^-8^) outside the HLA and surrounding 10Mb region for non-binary and binary phenotypes respectively.

Standing height was the trait with the largest number of hits (Figure 5) with 12,135 significantly associated variants distributed across 4,090 independent loci. We estimated that the leading polymorphisms across the 118 traits studied are distributed among 16,148 independent loci, therefore 25% of these independent loci contribute to the variation of height, as expected by a highly polygenic trait^4^.

**Figure 5.**
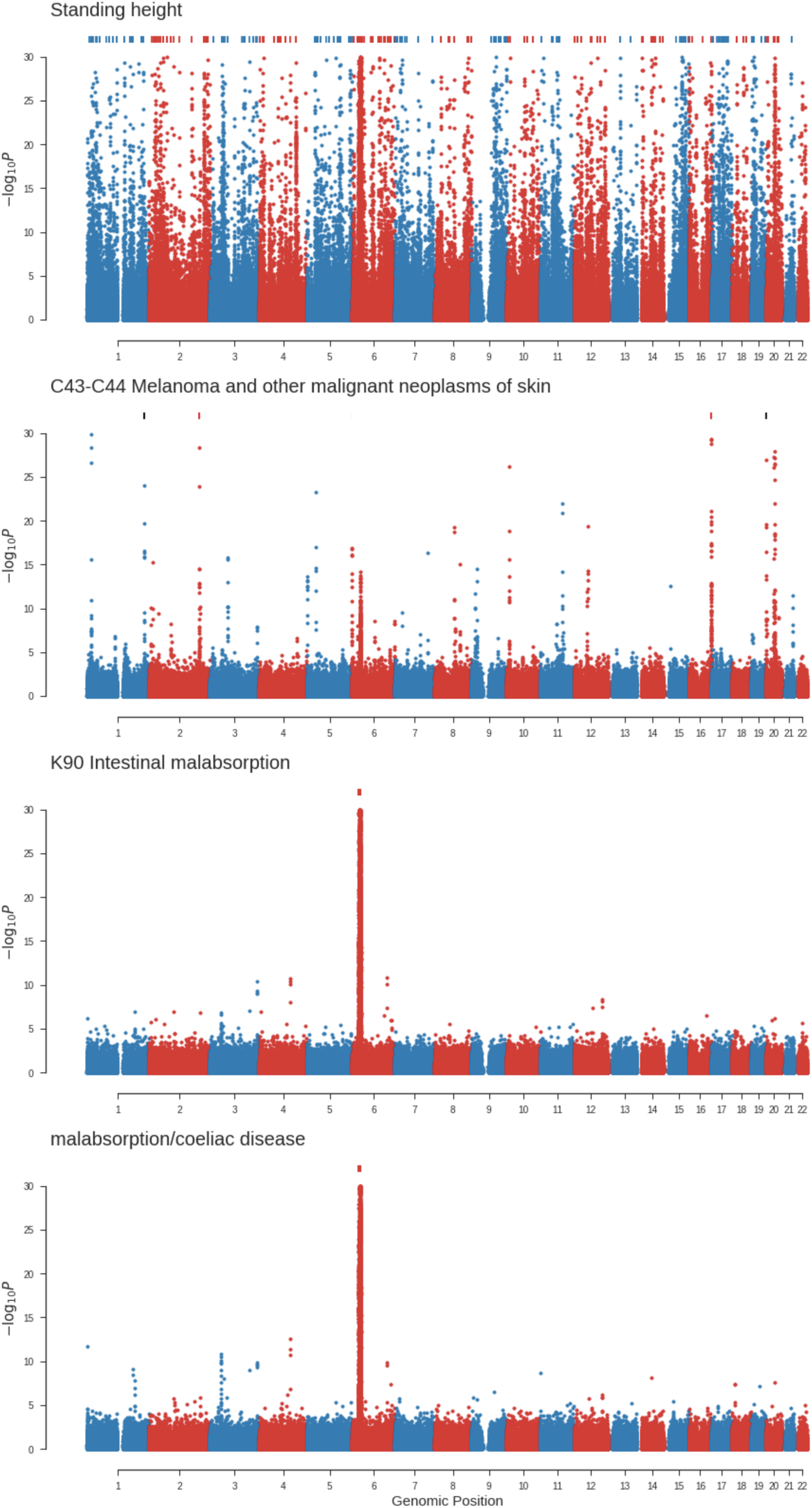
Manhattan plots for chosen phenotypes with the largest number of genome wide significant associations (P < 10^-8^) within each of four categories (all phenotypes, cancer registry, hospital episode statistics and self-reported non cancer illness). From top to bottom: amongst all phenotypes (Standing height), cancer registry phenotypes (Melanoma and other malignant neoplasms of skin), clinical information from hospital episode statistics and self-reported non cancer illness (intestinal malabsorption and malabsorption/coeliac disease respectively). Genetic variants with P < 10^-30^ are indicated by marks along the top of each plot.

### Distribution of genotyped GWAS hits among binary traits

The binary trait with the largest number of cases was self-reported hypertension, with an average across binary traits of 6,332 cases (Supplementary Table 1). Consistent with the reduced statistical power to detect association with binary phenotypes (mainly diseases) compared to non-binary traits we detected 49,974 associations at a P<10^-8^ (Supplementary Table 2), 80% of those were within the HLA region. One in ∼7500 genotyped variants was genome-wide significant (P<10^-8^) for binary traits, whilst 1 in ∼300 genotyped variants was significant for non-binary traits. Only genetic variants within the HLA region were associated with more than 20 binary traits each (Figure 3, Supplementary Fig. 3).

We found a positive correlation (r=0.83, P<10^-150^) between the heritability of the binary trait and the number of genome-wide significant variants, albeit of smaller magnitude to that found for the non-binary traits (Figure 4). Some of these traits were obvious outliers as they had large heritabilities but few significantly associated variants. The three largest heritabilities for binary traits were for three autoimmune diseases (ankylosing spondylitis, coeliac disease and seropositive rheumatoid arthritis) but few significant variants were found outside the HLA region. For instance, 1,031 out 1,035 genome-wide significant associations for ankylosing spondylitis were within the HLA region.

Among the categories for inclusion of genetic variants in the genotyping array there was a substantial enrichment for HLA (82%), ApoE (50%), Alzheimer's disease (43%) and cancer common variants (38%). The categories with the lowest enrichment were genome-wide coverage for low frequency variants (1.7%) and Rare, possibly disease causing, mutations (3.4%) (Supplementary Table 4).

We show three examples of Manhattan plots (Figure 5). The first example shows were there are associations with skin cancer (i.e melanoma and other malignant neoplasms of the skin). There are 329 variants associated (P<10^-8^) with skin cancer distributed among 88 independent loci. We found associations in genetic variants in or around known susceptibility genes (e.g. MC1R, IRF4, TERT, TYR) for melanoma^5^, but also novel associated genes like FOXP1 (rs13316357, P=1.6x10^-16^). FOXP1 has been reported to be a potential therapeutic target in cancer^6^ and its expression shown to change in a patient treated with immunotherapy^7^. The other two examples show the similarity between the results of one of the self-reported and clinically defined traits available in UK Biobank. The Manhattan plots for self-reported and clinically defined coeliac disease are very similar but not identical, which suggests that generally there will be benefit in analyzing both clinically and self-reported traits.

## DISCUSSION

We used circa 410,000 related and unrelated white-British UK Biobank participants to build the largest atlas of genetic associations to date. Summary statistics for 717 traits will be available to the research community to help them gain further insight into the genetic architecture of complex traits. Unlike other currently available databases, like the GWAS catalog (which contains 39,366 unique SNP-trait associations), our database includes significant and non-significant associations, providing thus an unbiased view of phenotype-genotype associations across a large number of traits. In addition, the database contains 76,345 independent genotype-phenotype associations, genetic and environmental correlations, and estimates of SNP heritability to allow researchers to perform their own filters on what a meaningful association or heritability is. We hope this database will be useful to those working on complex traits genetics, but also to those that have not got the expertise or capabilities to perform analyses at this scale.

## URLs

GeneATLAS, http://geneatlas.roslin.ed.ac.uk; UK Biobank, http://www.ukbiobank.ac.uk/; ARCHER UK National Supercomputing Service, http://www.archer.ac.uk; DISSECT, https://www.dissect.ed.ac.uk; GWAS catalog https://www.ebi.ac.uk/gwas/; Affymetrix array https://affymetrix.app.box.com/s/6gc2mcw2s6a7zbb7wijn

## ACKNOWLEDGEMENTS

This research has been conducted using the UK Biobank Resource under project 788. The work was funded by the Roslin Institute Strategic Programme Grant from the BBSRC (BB/P013732/1) and MRC grant (MR/N003179/1). AT also acknowledge funding from the Medical Research Council. Analyses were performed using the ARCHER UK National Supercomputing Service.

## AUTHOR CONTRIBUTIONS

All authors contributed equally to this work.

## COMPETING INTEREST STATEMENT

The authors declare no competing financial interests.

## ONLINE METHODS

### Phenotypes

In total we analysed 717 phenotypes in White-British UK Biobank participants. These included 596 binary phenotypes generated from self-reported disease status, ICD10 codes from hospitalization events, and ICD10 codes from cancer registry, as well as a further 3 binary and 118 non-binary (comprising continuous and integral measures) phenotypes from across the UK Biobank. A description of each phenotype, its category and the relevant UK Biobank fields can be found in Supplementary Table 1. Some of the traits analysed have some redundancy that has been left for completeness, that is some of these traits were measured in different ways during the study (e.g. weight) or are analysed as self-reported traits and clinical traits (e.g. malabsorption). For disease traits all individuals reporting a disease code were coded as cases with all other individuals considered controls. Only non-disease phenotypes with missing data rate < 5% were selected for analysis. For these phenotypes missing values were imputed to the age and sex specific mean in the White-British cohort.

### Genotypes

The genotypes of the UK Biobank participants were assayed using either of two genotyping arrays, the Affymetrix UK BiLEVE Axiom or Affymetrix UK Biobank Axiom array. These arrays were augmented by imputation of ∼96 million genetic variants from the Haplotype Reference Consortium^3^, the thousand genomes^8^ and the UK 10K^8^ projects. Full details regarding these data have been published elsewhere^9^.

We excluded individuals who were identified by the UK Biobank as outliers based on either genotyping missingness rate or heterogeneity, whose sex inferred from the genotypes did not match their self-reported sex and who were not of white British ancestry. Finally, we removed individuals with a missingness > 5% across variants which passed our QC procedure. The resulting White-British cohort comprised 408,455 individuals.

From the genotyped data we only retained bi-allelic autosomal variants which were assayed by both genotyping arrays employed by UK Biobank. We furthermore excluded variants which had failed UK Biobank quality control procedures in any of the genotyping batches. Additionally we excluded variants with P < 10^-50^ for departure from Hardy-Weinberg, computed on a subset of 344,057 unrelated (Kinship coefficient < 0.0442) individuals in the White-British cohort, and with a missingness rate > 2% in the White-British cohort. Only variants with MAF>10^-4^ in the White-British cohort were tested for association in the GWAS of the 717 traits, this cut-off corresponds to less than 82 occurrences of the minor allele in the White-British cohort.

### GWAS Analysis

To test each genetic variant whilst taking into account population structure in UK Biobank (e.g. presence of related individuals or local structure), we used a Linear Mixed Model. Specifically, the model takes the form

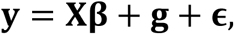

where **y** is the vector of phenotypes, **X**, is the matrix of fixed effects, and **β** the effect size of these effects. We included as fixed effects sex, array batch, UK Biobank Assessment Center, age, age^2^, and the leading 20 genomic principal components as computed by UK Biobank. **g** is the polygenic effect that captures the population structure, fitted as a random effect. It follows the distribution 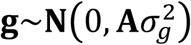, with **A** the Genomic Relationship Matrix (GRM), and 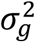 the variance explained by the additive genetic effects. The GRM was computed using common (MAF > 5%) genotyped variants that passed quality control. Finally, 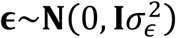 is a residual effect not accounted for by the fixed and random effects. Under this model, the phenotype vector **y**, follows the distribution 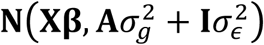.

Fitting one instance of such a LMM model is computationally very demanding. Following a naïve approach, the required computational time increasing with the cube of the sample size, ∼O(N^3^), and the memory requirements with the square of the sample size, ∼O(N^2^). Consequently, fitting a single model on a cohort of the size of UK Biobank is challenging, and fitting millions of these models, one for each analysed genetic variant and phenotype is not feasible with standard computational and statistical approaches. To address this problem, we took advantage of three different tools. First, we used a large supercomputer (5,040 processor cores working together for ∼10h, and using ∼5TB of memory for computing the GRM eigen-decomposition), and DISSECT^10^ to speed up the calculations. Second, we computed the full eigen decomposition of the GRM, **A** = Λ**Σ**Λ^*T*^, where Λ is the matrix of eigenvectors, and **Σ** is a diagonal matrix containing the eigenvalues. This allowed us to transform all the other model matrices, **y**, **X**, and **∊** to the new space where the GRM is diagonal. Although the eigen-decomposition is a computationally intensive process, once diagonalized, the computational time of fitting a model is reduced considerably to ∼O(N), thus enabling us to perform several tests using Mixed Linear Models on a cohort of hundreds of thousands of individuals. Finally we performed over 23 billion tests using a two-step approximation that optimizes the computational resources^11^. The first step of the approximation fits a LMM that adjusts by the relevant fix (e.g. age, sex, etc.) and random effects (genetic effects) to each trait, the second step uses the residuals of LMM to test all available genetic markers for significance in a linear model. We adjusted for the genetic variants genotyped in the odd chromosomes when testing polymorphisms in the even chromosomes, and for the genetic variants genotyped in the even chromosomes when testing genetic variants in the odd chromosomes.

### HLA Region

We defined the *HLA* region as the region of chromosome 6 spanning base pairs 28,866,528 to 33,775,446. Throughout all analyses we included 10Mb either side of the above *HLA* region to account for LD with variants outside this region.

### Estimation of Genetic Parameters

In order to estimate heritabilities and genetic correlations we fitted LMMs for each trait with a GRM containing all common (MAF > 5%) autosomal genetic variants which passed QC. The heritability was estimated as 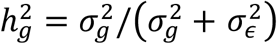, where 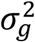 and 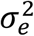 are the estimates of the genetic and residual variance. For all binary outcomes, we transformed heritabilities on the observed scaled to the liability scale using the population prevalence of the disease. We provide sex-specific prevalences to allow sex-specific transformations (Supplementary Table 1). Using the model fits we computed best linear unbiased predictor estimates of genetic additive values for each individual. The genetic correlations were estimated by computing correlations between these additive genetic values. Environmental correlations were estimated as 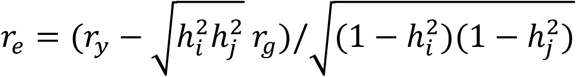 where *r*_*y*_, *r*_*g*_ are the phenotypic and genetic correlations for traits *i*, *j* and.

### Independent Loci

We clustered GWAS results into independent loci using the –`clump` option of the plink 1.9 software^12^. Specifically for each trait individually, we clustered GWAS results by selecting genome wide significant variants as lead variants and assigning to them unassigned variants within 10Mb, that have P<10^-2^ and a *r*^2^ > 0.3 with the lead variant. To compute the total number of independent loci across all traits, we performed the same clustering on the lead variants of loci across all traits, choosing the lowest P value for variants which were lead variants of a locus in different traits.

### Relation of association count and chromosome length

We regressed the number of significant associations (P<10^-8^) across traits for each chromosome on the covered length of the chromosome, i.e., distance in base pairs of the first and last genetic variants contained on the genotyping array, and the number of genetic variants on the chromosome contained on the genotyping array. For chromosome 6 we excluded the HLA region and variants contained therein from the statistics. We compared the full model to one with either the chromosomal length or number of genetic variants removed using the likelihood ratio test. The full model was not significantly better than either of the reduced models, which both were significant when compared to a null model containing only an intercept.

